# cMYC protein interactions and chromatin association in NUT carcinoma

**DOI:** 10.1101/2022.04.22.489219

**Authors:** Barry M. Zee, Artyom A. Alekseyenko, Anne E. Smolko, Hyuckjoon Kang, Mitzi I. Kuroda

## Abstract

The oncogene c*MYC* (HGNC:7553) is a critical genomic target of the BRD4-NUT (B4N) protein that defines many of the NUTM1-rearrangement cancer subtypes in NUT carcinoma (NC). Previously, we reported that B4N interacts with the EP300 lysine acetyltransferase (KAT3B) and creates hyperacetylated “megadomains” that activate downstream genes such as *cMYC*. Here we ask how misregulated cMYC in turn interacts with protein partners and target genes in patient-derived NC797 cells, and whether these interactions change in response to B4N inactivation. We used CRISPR-Cas9 mediated knock-in of a BioTAP affinity tag to analyze cMYC protein expressed from its normal chromosomal context. This allowed us to implement a crosslinking purification method termed BioTAP-XL to preserve cMYC integrity and chromatin association for genomic and mass spectrometry-based proteomic analyses. We found that in the NC797 cell line, cMYC interacts primarily with the NuA4 KAT5 lysine acetyltransferase complex, with interactions that are maintained despite a decrease in cMYC levels after JQ1 treatment. We propose that a cascade of aberrant acetyltransferase activities in NC797 cells, first via EP300 recruitment by B4N to mis-regulate *cMYC*, and then by KAT5 interaction with the resulting overexpressed cMYC protein, drive NC cell proliferation and blockade to differentiation.

**Graphical Abstract:** 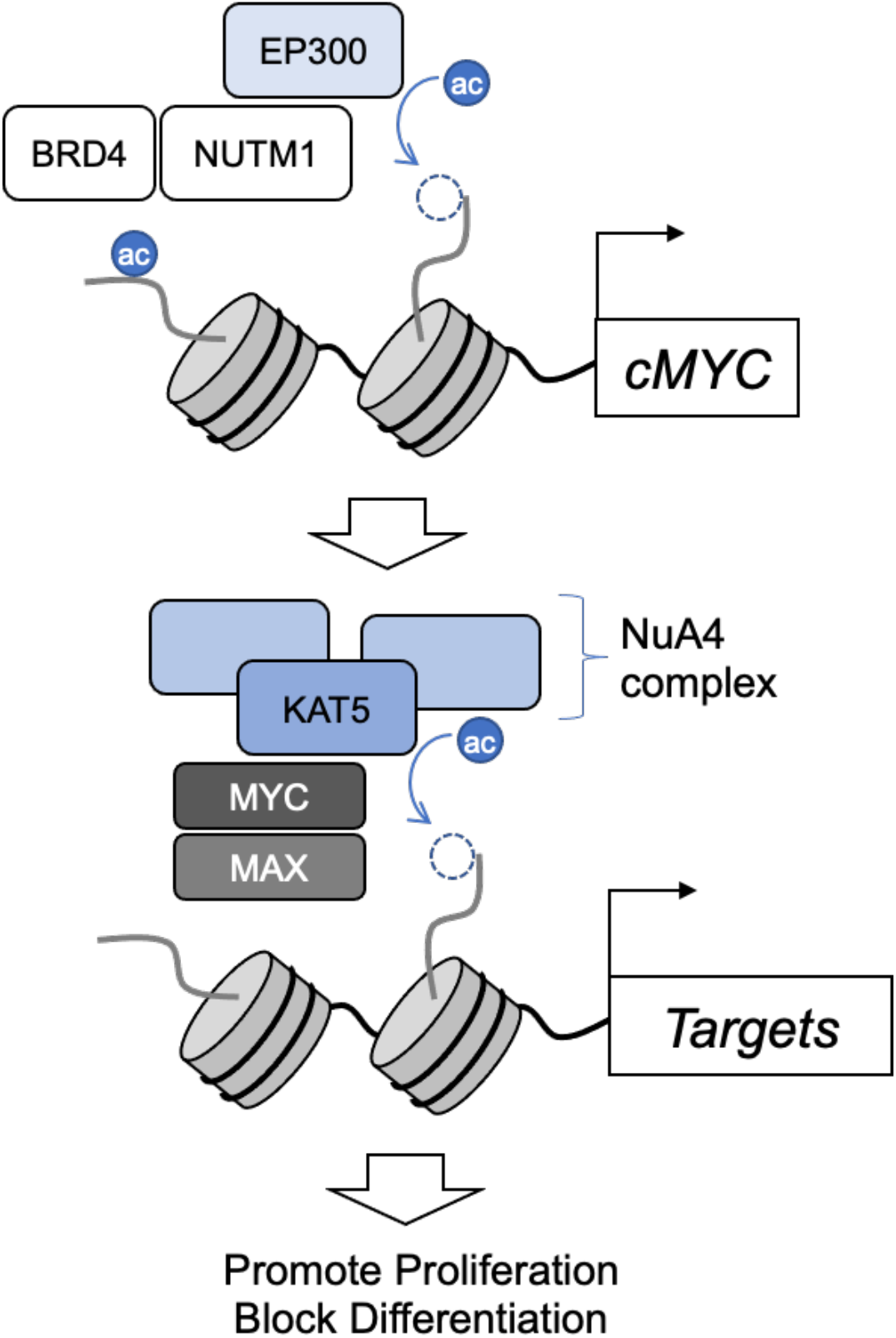

**Simple Summary:** A longstanding goal in biology is to understand how protein interactions influence and reflect cellular disease states. MYC is a critical regulator of cellular proliferation that is mis-regulated in many cancers, including those with NUTM1-rearrangements featured in this issue. NUT carcinoma cells are dependent on MYC expression, which blocks differentiation. Using a crosslinking approach to identify MYC protein interactions in a NUTM1-rearranged patient cell line, we found that MYC interacts primarily with subunits of the NuA4 lysine acetyltransferase (KAT5), one of two KAT complexes previously discovered as MYC interactors in non-NUTM1 cell lines.

## 1. Introduction

NUT-rearrangement cancers are a collection of aggressive and poorly-differentiated squamous cell carcinomas whose unique molecular hallmark is somatic expression of *NUTM1* (HGNC:29919)[1]. Chromosomal rearrangements result in the otherwise germline-restricted *NUTM1* gene to be fused downstream of a somatically expressed gene, most commonly *BRD4* (HGNC:13575) to generate the fusion protein BRD4-NUT (B4N). At least two NC fusion proteins: B4N and ZNF532-NUT (a fusion product with *ZNF532* (HGNC:30940)) have been shown to regulate expression of the proto-oncogene *cMYC* (hereafter referred to as *MYC*). This is thought to occur through an aberrant, feed-forward loop in which the NUTM1 portion of the fusion protein attracts the EP300 lysine acetyltransferase, while the BRD4 bromodomain binds the resulting acetylated nucleosomes. The resulting formation of unusually large, hyperacetylated chromatin regions termed “megadomains” drives increased expression of underlying genes and regulatory regions [2, 3]. Treatment of NC patient cells in culture with a small molecule, JQ1, that inhibits interaction of the BRD4 bromodomains with acetylated lysines, leads to loss of megadomains, down-regulation of *MYC*, and differentiation of B4N cells [2, 4]. The unique importance of *MYC* was revealed in studies where induced *MYC* transgene expression reversed the differentiation induction caused by B4N knockdown [5]. Furthermore, non-B4N-dependent pathways such as *KLF4* (HGNC:6348) expression help maintain *MYC* levels in cells resistant to JQ1 [6]. Thus, understanding the mechanism of action of MYC in NC cells should provide valuable insights into this therapeutically intractable disease.

Much of what is understood about MYC stems from non-NC cells [7-13]. MYC binds DNA as a heterodimer with MAX, a second bHLHZip (basic helix-loop-helix leucine zipper) protein [14] and positively regulates the expression of critical enzymes involved in growth and metabolism, thereby increasing global RNA and protein expression [9]. Although the direct target specificity of MYC has been a subject of considerable debate, this is likely due to an incomplete picture of the relationship between binding and function. MYC can be observed to bind to most transcription start sites (TSS), enhancers, and accessible chromatin in various cell types, as it has relatively strong non-specific DNA binding in addition to its sequence-specificity for E-box motifs [15, 16]. Current genetic and functional analyses are consistent with direct regulation by MYC of genes involved in nucleotide biosynthesis, RNA metabolism, and ribosome biogenesis, which in turn drive numerous secondary changes in gene expression [10, 16].

In addition to heterodimerization with MAX, early studies identified MYC protein interactions with the SAGA/STAGA and NuA4 histone acetyltransferase complexes [17-22], linked to MYC via the TRRAP adaptor protein [23, 24]. These multi-subunit complexes [25, 26] each function as transcriptional coactivators, with enzymatic activities targeted mainly to histone H3 in the case of SAGA/ STAGA, and H2A, H2AZ, and H4 in the case of NuA4 [24].

For many years comprehensive proteomic analyses of MYC interactions were impeded by technical challenges. First, in normal cells the MYC protein has a relatively short half-life of approximately 30 minutes, a result of ubiquitin-dependent and ubiquitin-independent pathways and of the even shorter (10 minute) half-life of *MYC* RNA itself [27-30]. Second, expression of the *MYC* gene depends on many long-range distal regulatory elements [31-34] suggesting the importance of complex regulatory circuits of the *MYC* locus which ideally should be preserved when analyzing MYC in different cell states. Finally, it has been difficult to extract intact MYC complexes from chromatin using traditional biochemical methods. This technical challenge has been overcome by proximity-dependent biotin identification (BioID) with considerable success [12, 35]. Here, we apply an orthogonal approach, BioTAP-XL [3, 36], to characterize MYC protein and genomic interactions in NC and non-NC cells. BioTAP-XL employs crosslinking, stringent two-step affinity purification, and analysis of enrichment over input, thus diminishing background and potentially allowing the most significant interactions to rise to the top. Consistent with the previous studies [12, 17-22, 35, 37], we found that subunits of the NuA4 and STAGA lysine acetytransferases were top MYC interactors in a non-NC cell line. In contrast, NuA4 was the predominant MYC interactor in an NC cell line, and these interactions were largely maintained after JQ1 treatment reduced overall MYC levels, suggesting that protein complex assembly may be independent of differentiation status. As chromatin acetylation complexes were the most enriched MYC interactors using our stringent cross-linking, tandem affinity purification approach, our results support models in which acetylation is likely to be central to the biological function of MYC.

## 2. Materials and Methods

### 2.1. Generation of N-BioTAP-cMYC NC797 cells using CRISPR/Cas9

To generate N-BioTAP and N-ProteinA (PrA) cMYC fusions in NC797 cells [38], the following DNA fragments: pAVV-MCS-NheI-5’cMyc_BamHI-ATG-loxP-Blasti-P2A-loxP-PrA-Bio-3’cMyc and pAAV-5’-cMYC-ATG-LoxP-Blasti-P2A-loxP-PrA-3’-cMYC respectively, were synthesized as gBlock gene fragments (Integrated DNA Technologies, **Datafile S1**) and introduced into the NheI/BamHI restriction sites of pAAV-MCS2 by Gibson assembly (New England Biolabs). N-BioTAP and N-PrA cassettes targeted the first ATG of the *MYC* gene. The pAAV-nEFCas9 (Addgene, plasmid 87115) was used to express Cas9, and the pAAV-tagBFP U6-gRNA expression vector was used to express a gRNA targeting the start codon of cMYC: gRNA-cMYC-start (**Table 1**). Recombinant AAV2 was packaged in HEK293T cells using pHelper and pRC2-mi342 plasmids (Takara, Catalog #632608). Three days after transfection, cells were harvested and AAV2 was isolated using AAVpro Extraction Solution (Takara, Catalog #6235). Two million NC797 cells were infected with 30ul of each adeno-associated virus (AAV2). Following 1 week of culture post-AAV infection, cells were selected with 7.5ug/ml Blasticidin (Invitrogen). Surviving cells were re-plated and single-cell clones were isolated and expanded, and genotyping PCR and sequencing were performed to check for proper integration of the BioTAP or PrA cassette in the c-MYC locus. We recovered both heterozygous and homozygous clones. Homozygosity of the cassette insertion was determined by absence of the wildtype PCR fragment (1033bp) and presence of the larger PCR fragment (2083bp) with the following PCR primers: Genom-Myc(1S) and Genom-Myc(1A) (**Table 1**). Next, we validated the presence of the correctly sized tagged MYC protein and absence of wild type MYC by Western blot probed with a c-MYC antibody (Rabbit mAb, Cell Signaling Technology, Cat# 5605). We selected homozygous clone number 2 (for N-BioTAP-cMYC) and clone number 4 (for N-PrA-cMYC) for Cre-dependent cassette excision experiments.

**Table 1.**
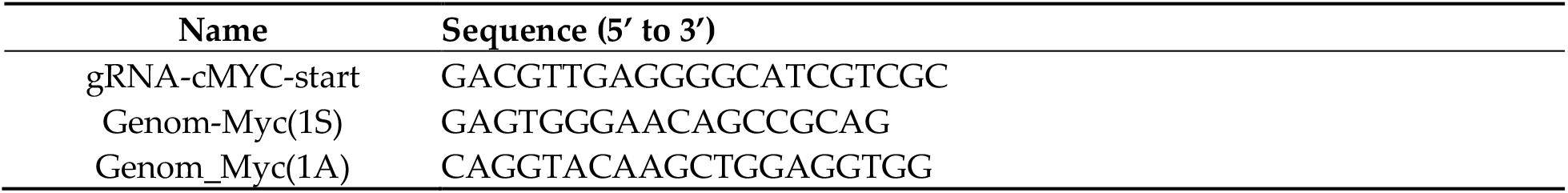
Sequences of oligonucleotides used in paper.

Next, the recombinant pAAV-CRE plasmid was packaged in HEK293T cells using pHelper and pRC2-mi342 plasmids (Takara, Catalog #632608). Cells were harvested three days after transfection and AAV2 was isolated using AAVpro Extraction Solution (Takara, Catalog #6235). Two million 797 cells with homozygous insertions (clones 2 or 4) were infected with 30ul of each adeno-associated CRE virus (AAV2). Following one week of culture post-AAV infection, cells were re-plated, single-cell clones were isolated and expanded, and genotyping PCR and sequencing were performed to check for absence of the drug resistance cassette in the c-MYC locus. We selected clone number 2-7 (for N-BioTAP-cMYC) and clone number 4-6 (for N-PrA-cMYC) to proceed with LC-MS and ChIP-seq experiments.

### 2.2. Generation of inducible N and C-BioTAP-cMYC expressing HEK293T cells

We used lentiviral transduction to generate stable HEK293T cell lines [2] carrying cMYC constructs with the BioTAP tag at either the N or the C terminus to induce transcription of these clones from the CMV/2xtetO promoter for genetic complementation experiments. To make N- and C-terminal BioTAP constructs, we used the Gateway recombination system to introduce the following Myc ORF clone HsCD00039771 in pDONR221from the CCSB Human ORFeome Collection into pHAGE-TRE-DEST-NBioTAP (Addgene #53568) and pHAGE-TRE-DEST-CBioTAP lentiviral vectors (Addgene #53569). Stable 293T-Trex-cell lines containing each transgene were generated by lentiviral transduction in the presence of 8μg/mL polybrene, followed by selection with 2μg/mL puromycin dihydrochloride (Sigma #P8833) as described previously [39]. To induce transcription of the cDNA clones from the CMV/2xtetO promoter we added doxycycline (1μg/mL) to the medium for 48 hours.

### 2.3. Proteomic analysis of cMYC interactions using BioTAP-XL

All cell lines were cultured as monolayers in DMEM (Invitrogen) supplemented with 1x Penicillin Streptomycin (Hyclone, South Logan, UT), 1x Glutamax (Gibco), and 10% (v/v) Fetal bovine serum (FBS) (Hyclone). For JQ1 treatment, 500uM of JQ1 (gift from Dr. C.A. French) was added to NC797 cells for various durations before harvesting for Western Blot analysis. The 12 hour timepoint was ultimately chosen for BioTAP-XL experiments.

Our primary experiments were performed with 2-7 N-BioTAP-cMYC (clone 2-7). 2.0 × 10^9 cells were grown as monolayer cultures in 150-mm dishes. 100 plates total were used for BioTAP-XL purification. The main steps of the BioTAP-XL procedure were as previously described [39]. For single-step BioTAP-XL, proteins enriched by IgG-agarose beads were eluted and processed with Agencourt AMPure-XP magnetic beads (Beckman Coulter) as previously described [40]. In brief, fifty micrograms of eluted proteins were acidified and diluted with acetonitrile prior to binding by magnetic beads. SDS was removed with ethanol and acetonitrile washes, and proteins were digested with overnight trypsin incubation, yielding peptides for desalting and LC-MS analysis. LC-MS acquisition and analyses were performed as previously described [3]. Offline peptide fractionation was not performed. Rather, peptide samples were autosampler loaded onto a 100µm ID fused silica capillary column packed with 25cm of 2.6µm Accucore C18-resin (Thermo Fisher) and electrosprayed into either a Velos-Orbitrap Pro or Velos-Orbitrap Elite mass spectrometer (Thermo Fisher). Peptides were resolved with a gradient from 0% to 5% Buffer B from 0 to 10 minutes, from 5% to 32% Buffer B from 10 to 65 minutes, and from 32% to 95% Buffer B from 65 to 68 minutes at a flow rate of approximately 110nl/min delivered on an Accela pump (Thermo Fisher), where Buffer A composition is 97.5% water, 2.4% acetonitrile, 0.1% formic acid and Buffer B composition is 7.4% water, 92.5% acetonitrile, 0.1% formic acid. Peptides were analyzed under Top22 data-dependent mode with collision-induced dissociation (CID), using the Orbitrap detector for MS1 acquisition and ion-trap detector for MS2 acquisition. Peptides were identified with SEQUEST against the human proteome with precursor mass tolerance at 20ppm and fragment ion mass tolerance at 0.9 Da. Methionine oxidation was set as dynamic modification. Identifications were further filtered at a 2% false discovery rate. The number of total spectra number for each protein were used to determine relative abundance. Up to two technical replicates per input or immunoprecipitated samples were acquired.

### 2.3. Genomic analysis of cMYC binding

ChIP-seq analysis was performed in parallel with proteomic analyses as previously described [36]. Raw fastq files were aligned to the human genome (hg38) using bowtie2.3.4.3 with default settings [41]. Duplicate reads were then removed with samtools1.14 [42]. Peak calling was performed on DMSO treated samples with HOMER4.9 using the findPeaks command [43]. HOMER annotatePeaks.pl was used to assign peaks to genomic regions. Deeptools3.0.2 was used to generate bigwig files, enrichment profiles, and heatmaps [44]. Bigwig files were normalized to 1x genome coverage. For heatmaps and enrichment profiles, MYC signal was normalized to its corresponding input and enrichment was shown as log_2_(IP/input). Differential peak analysis was performed with the DiffBind package in RStudio using default settings: (http://bioconductor.org/packages/release/bioc/vignettes/DiffBind/inst/doc/DiffBind.pdf).Datasets can be obtained from the NCBI GEO database (accession# GSE201033).

## 3. Results

### 3.1. BioTAP-XL strategy for analysis of MYC interactions

To address the potential instability or loss of MYC during conventional biochemical extraction from DNA, we applied a crosslinking approach termed BioTAP-XL for MYC complex purification in NC797 cells, an NC patient-derived cell line, and in HEK293 cells, a non-NC line for comparison. The BioTAP affinity tag itself is composed of the Z domain from *Staphylococcal* protein A and a biotinylation target domain from *Propionibacterium* transcarboxylase. Biotinylation of this target sequence occurs in most organisms without requiring introduction of an exogenous biotin ligase or addition of biotin to the media [36]. After generating a cell line that expresses the BioTAP-tagged bait, the BioTAP-XL workflow involves three steps: 1) formaldehyde to inactivate cellular enzymes and to crosslink proximal protein and nucleic acid interactions, 2) sonication to solubilize the crosslinked chromatin pellet, and 3) enrichment of the tagged bait using tandem ProteinA and streptavidin pulldowns combined with stringent washes to reduce nonspecific interactions. This approach has successfully characterized the proteomic and genomic interactions of B4N, wild-type BRD4, ZNF532-NUT, wild-type ZNF532, and other proteins [3].

### 3.2. NuA4 lysine acetyltransferase subunits are highly enriched in BioTAP-MYC purifications

To characterize the protein interactions of N-BioTAP-MYC in NC797 cells, we used a CRISPR-Cas9 mediated knock-in approach. We successfully inserted the BioTAP affinity tag into both copies of the endogenous *cMYC* gene in NC797 cells (see Methods, **Figure 1**). A critical concern with interpreting affinity tag results is whether the tagged protein preserves the functions of the non-tagged protein. In NC models such as NC797 cells, B4N regulates transcription of the *MYC* gene, and NC797 cells are dependent on functional MYC expression [1, 2, 4, 5]. Since knockdown of MYC was previously shown to induce differentiation of NMC cells [5], that we could create and grow homozygous knock-in NC797 cells in large numbers (>10^9^) strongly suggests that the epitope tags do not interfere with normal MYC function.

**Figure 1.**
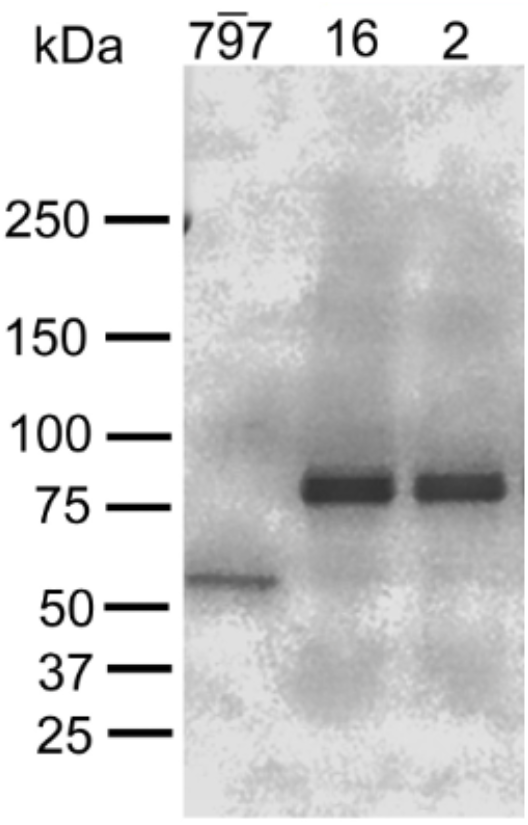
Western blot of NC797 cells carrying homozygous knock-in alleles of BioTAP-tagged MYC probed with anti-MYC (D84C12 Rabbit mAb, Cell Signalling Technologies). The untagged size of MYC in unedited NC797 cells is ~57 Kd, (lane 1). The size of N-BioTAP-MYC is ~80 Kd, seen in Clones 16 and 2 (lanes 2 and 3). Clone 2 was used for subsequent experiments. The original gel image can be viewed in Supplemental Figure 1.

We performed BioTAP-XL to enrich for MYC containing protein complexes and used liquid chromatography-mass spectrometry (LC-MS) to sequence the peptides resulting from trypsin digestion of the enriched complexes. MYC protein interactions were identified as the proteins with the greatest fold enrichment in total peptides identified in the pulldown compared to the input sample (**Datafile S2**). Fold-enrichment was calculated as the number of total peptides mapping to each protein divided by the molecular weight of the protein as a mass-normalized ratio, which was subsequently divided by the sum of all the mass-normalized ratios across all proteins identified in a given LCMS sample, and finally, compared between samples as log ratios.

We observed consistency (Pearson > 0.8) in overall enrichment of proteins between technical replicates of our BioTAP-XL experiments (**Supplemental Figure 2**). As expected, we recovered both MYC and its heterodimerization partner MAX [14] as among the top one to two percentile most enriched proteins (corresponding in this case to approximately twenty-three proteins). Beyond MYC and MAX, eleven of the most enriched proteins are related to chromatin function. In particular, eight are components of the NuA4 histone acetyltransferase complex [26, 37] (**Figure 2**). The enriched components include DMAP1 (HGNC:18291), EP400 (HGNC:11958), EPC1 (HGNC:19876), EPC2 (HGNC:24543), KAT5 (HGNC:5275), MBTD1 (HGNC:19866), TRRAP (HGNC:12347), and YEATS4 (HGNC:24859).

**Figure 2.**
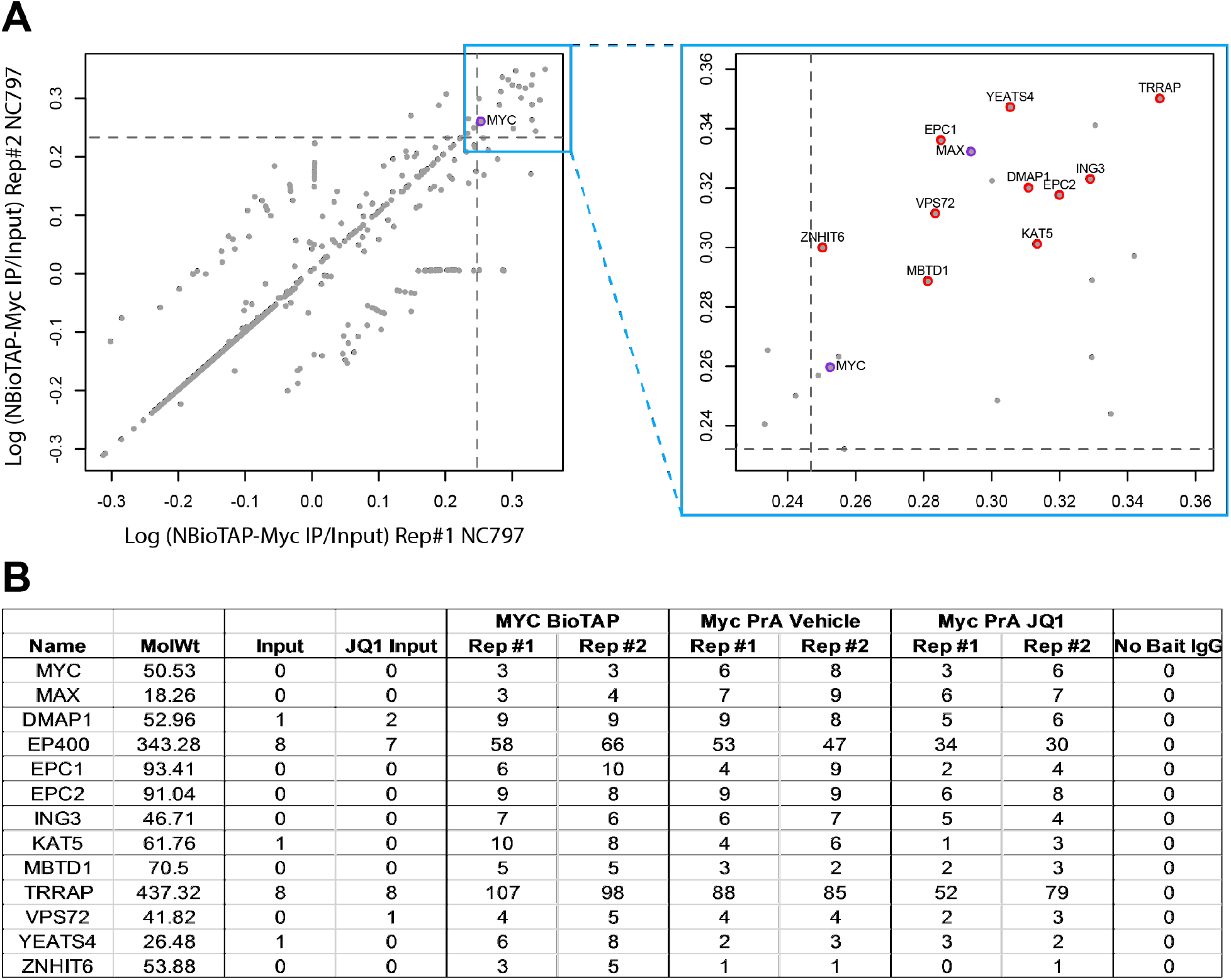
Characterization of MYC protein interactions in NC797 cells. (a) Scatterplots depicting enrichment of proteins from N-terminally tagged MYC using BioTAP-XL workflow in NC797 cells compared to non-enriched input. Each gray dot represents an individually identified protein, where its position is a measure of enrichment efficiency based on the number of total peptides identified in pulldown compared to input and normalized to the molecular weight of the protein from two technical replicates. The dashed gray lines denote the 98^th^ percentile of enrichment. The right plot is a zoomed in region of the most enriched proteins in the left plot from replicate experiments. (b) Number of total peptides mapping to each protein from a given pulldown experiment in independent (BioTAP-vs. PrA-tagged) NC797 cell isolates.

### 3.3. Comparison of MYC interactions in distinct cell types and cell states

To test whether the enrichment of NuA4 but not STAGA might be related to our crosslinking method rather than the NC cell type, we performed the BioTAP-XL experiment in the non-NC cell line HEK293-TREx, which was used in previous studies [12]. Similar to NC797 cells, we found enrichment of NuA4 components with the MYC-BioTAP bait in HEK293-TREx cells, but we also observed enrichment of SAGA/STAGA components such as ATXN7 (HGNC:10560) and KAT2A (HGNC:4201) (**Supplemental Figure 3, Datafile S3**). Therefore, MYC interaction with NuA4 but not STAGA appears to be a characteristic of the NC797 cell line rather than somehow biased by our crosslinking approach.

To test whether the NuA4 interaction with MYC would change upon B4N inactivation and the resulting cell state transition towards differentiation, we performed affinity enrichment of MYC after cells were treated with the bromodomain inhibitor JQ1. We knew from prior experience that the amount of MYC bait protein expression after inactivation of B4N would be significantly decreased, which might be insufficient for a full tandem affinity purification. Therefore we modified our approach to utilize a single rather than dual tag, creating NC797 clones in which the endogenous gene had only the ProteinA portion of the affinity tag (PrA-MYC). Consistent with B4N directly regulating the endogenous *MYC* locus, treatment of NC797 cells with JQ1 reduced levels of PrA-MYC in a time-dependent manner (**Figure 3**). That cells relying on epitope-tagged-MYC still responded to JQ1 provided further evidence for *MYC* functionality after tag insertion. Although we had hoped to uncover regulation of MYC at the level of protein-protein interactions, we found that NuA4 interactions with MYC were largely preserved after a 12 hour treatment with JQ1 corresponding to a five-fold reduction in PrA-MYC levels (**Figure 2B**). To test whether recovery of NuA4 components could be due to non-specific capture of the proteins by the IgG agarose beads, we performed a parallel pulldown with NC797 cells expressing no tagged bait. We observed no recovery of peptides from MYC or NuA4 components (**Figure 2B**). Taken together, our results confirm the strong relationship between MYC and NuA4 in NC797 cells.

**Figure 3.**
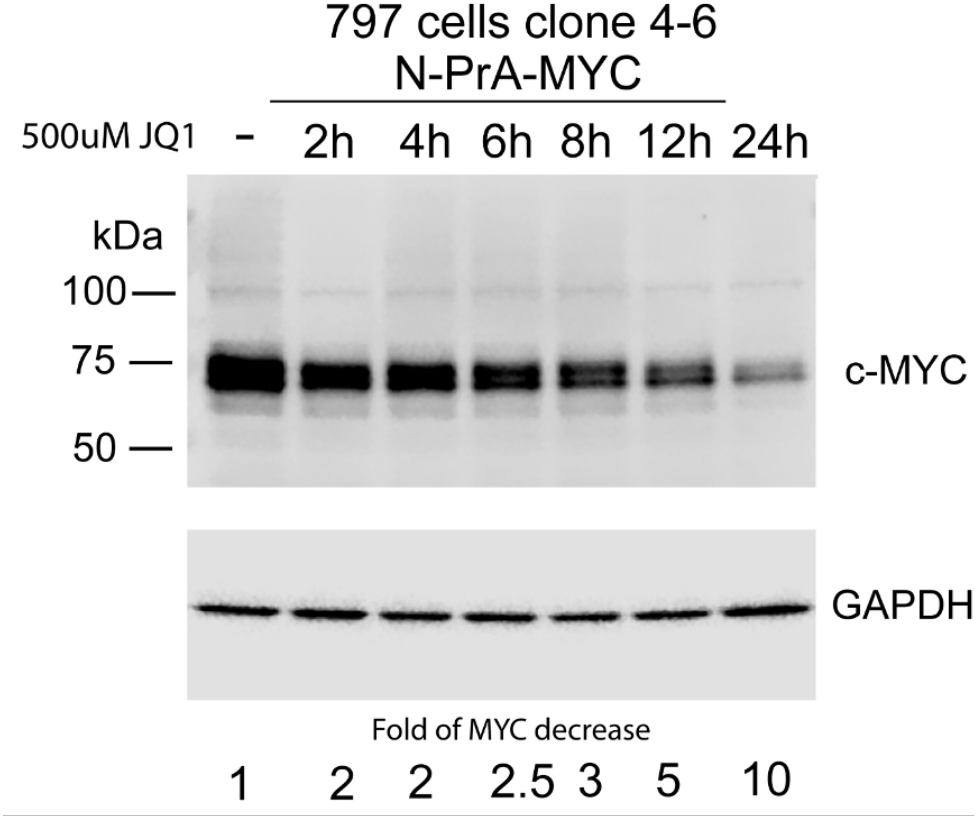
Effect of JQ1 treatment on MYC protein expression. Levels of MYC protein in N-PrA-cMyc-NC797 cells were assayed by Western blot, following different durations of 500uM JQ1 treatment, with approximate fold change in MYC levels provided below the blot compared to no treatment. The original gel image can be seen in Supplemental Figure S1.

### 3.4. Genomic profiling confirms broad MYC association largely unchanged by JQ1 treatment

In parallel with the proteomic analyses, we mapped the genomic occupancy of PrA-MYC in NC797 cells following JQ1 or DMSO treatment. Differential binding analysis resulted in 2,238 regions with significantly altered MYC binding (FDR < 0.05) (**Figure 4A,B**). Consistent with the decrease in MYC levels, all but 4 of these MYC-bound regions showed decreased occupancy(n=2,234). Significantly downregulated regions were located at promoters, defined as +/-1 Kb from transcription start sites (25%, n=555/2,234), introns (40%, n=900/2,234), and intergenic regions (28%, n=623/2,234) (**Figure 4C)**. While MYC occupancy was diminished, the identities of MYC binding sites were not globally changed by JQ1 treatment (**Figure 4B,D**). It is known that JQ1 treatment alters gene expression [1, 2], but in retrospect, the lack of overall change in genomic occupancy is not surprising given the recent discovery that pervasive non-specific DNA binding contributes to, but also obscures the considerably smaller set of specific regulatory targets of MYC [15, 16]. In summary, our comparison of proliferating and differentiating NC cells suggests that MYC-dependent changes in cell state generally occur without altering the identity of the most enriched MYC protein interactions or overall DNA binding sites.

**Figure 4.**
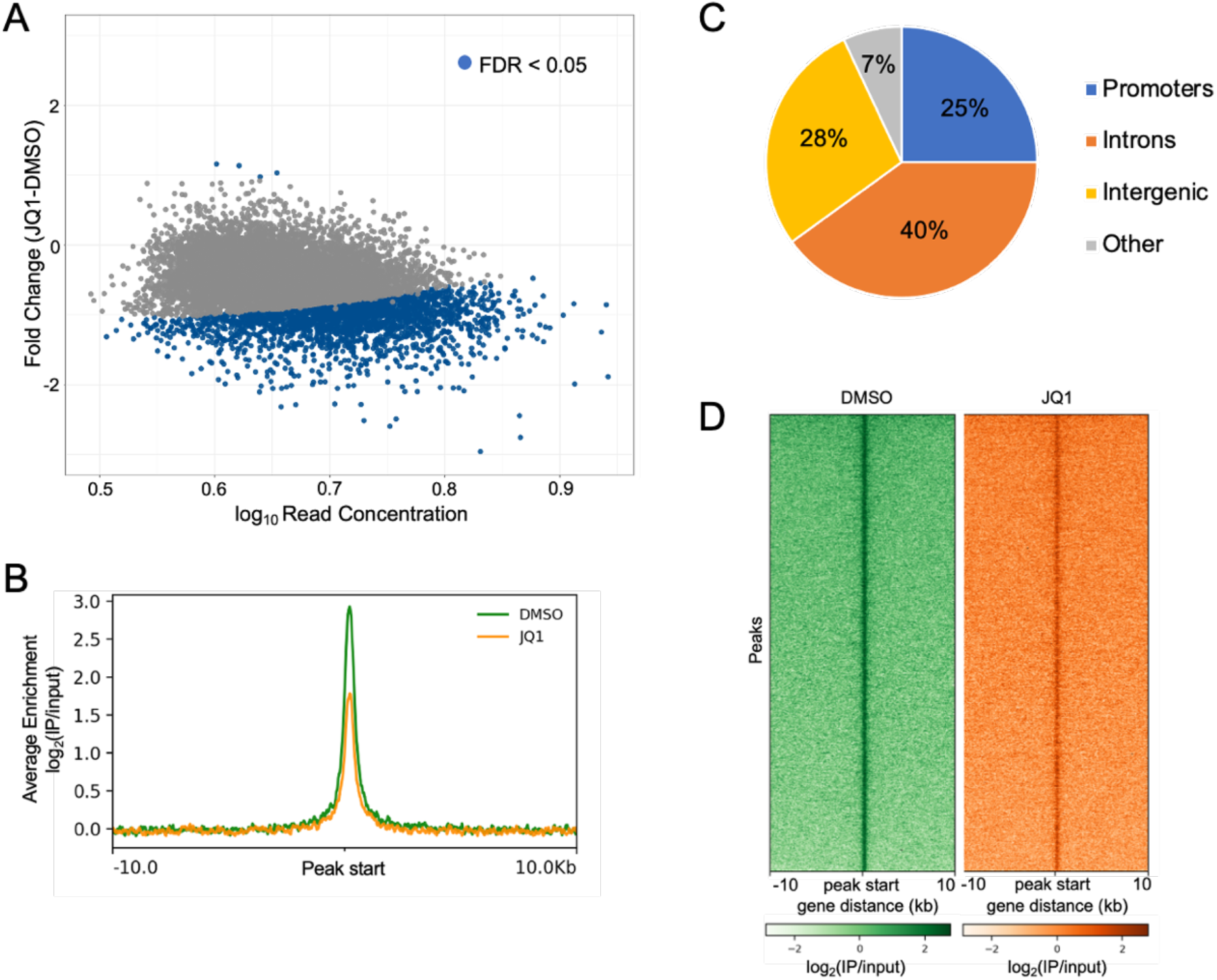
Effect of JQ1 on MYC genomic binding. (a.) MA plot of MYC differential binding after JQ1 treatment. Each point represents a 400 bp MYC-bound region. Blue points indicate significantly altered regions (FDR < 0.05). The x-axis depicts the log_10_ of the average read concentration across all samples. The y-axis indicates the fold change after JQ1 treatment. (b.) Enrichment profile plotting the average signal of significantly decreased MYC-bound regions in DMSO and JQ1 treated samples. One representative replicate is shown. (c.) Pie chart depicting the percentage of significantly decreased MYC-bound regions mapped to the specified genomic regions. (d.) Heatmap of MYC enrichment at significantly decreased MYC-bound regions in DMSO and JQ1 treated samples. One representative replicate is shown.

## 4. Discussion

To characterize MYC protein partners in NC797 cells, we applied a crosslinking affinity pulldown method. We took this approach because, historically, MYC complexes have been difficult to purify. However, significant progress has been made in this regard through recent BioID studies by the Penn laboratory, in which proximity labeling resulted in a comprehensive study of MYC interactors [12]. Extending the known association with acetyltransferases [17-22], they mapped the interactions of both NuA4 and STAGA complexes to MYC homology box II (MBII), showed that MBII is linked to both complexes through TRRAP, and confirmed the importance of MBII in stimulation of acetylation by MYC. Our data in HEK293T cells using the orthogonal BioTAP-XL approach is in agreement with the association of MYC with both NuA4 and STAGA. It remains to be determined why the NC cells we studied here show strong enrichment for NuA4 but not STAGA. In this regard, we hope that our datasets will be useful in further study of MYC and downstream targets specifically in NC cells. Demonstrations that MYC is dependent on acetyltransferases for its function have suggested potential therapeutic options to what is generally considered an undruggable oncoprotein [45, 46]. Our results suggest that development of specific inhibitors of KAT5 for synergy with JQ1 could be particularly relevant to the treatment of NC.

## 5. Conclusion

MYC is a central player in a majority of cancers, yet understanding its regulation, targeting, and mode of action are important unsolved problems. When we initiated this study, we knew that the B4N oncogenic fusion attracted the p300 acetyltransferase to drive oncogenesis, with the *MYC* gene as a critical target [1]. Since MYC function in NC cells is reversible by the JQ1 small molecule BET bromodomain inhibitor, we saw this as an opportunity to compare MYC interactions in the same cells under oncogenic and differentiating conditions. While we did not find a difference in protein interactions or gene targeting in the distinct growth states, we identified a close association between MYC and the NuA4 acetyltransferase in NC cells. We propose that future analysis of MYC transcriptional regulation in NUT carcinoma, and likely other cancers, should prioritize how MYC recruits KAT complexes to modify the chromatin of its many gene targets.

## Supporting information

Datafile S2

Datafile S3

Datafile S1

## Supplementary Materials

Fig. S1: Original gel images for Figs. 1 and 3.

Fig. S2. Correlation of NC797 BioTAP-XL replicates.

Fig. S3. Comparison of MYC interactions with NuA4 and SAGA/STAGA components in HEK293 and NC797 cells.

Datafile S1: DNA sequences used to create NC797 cells with N-terminal fusions of the BioTAP- or Protein A-epitope tags to the endogenous cMYC coding region.

Datafile S2: Total peptide counts of NC797 proteomic results.

Datafile S3: Total peptide counts of HEK293T proteomic results.

## Author Contributions

Conceptualization, AAA and MIK; methodology, validation, formal analysis, investigation, BMZ and AAA; resources, data curation, BMZ, AES, and HJK; writing – original draft preparation BMZ and AAA; writing – review and editing, BMZ and MIK; visualization, BMZ, AAA, AES and HJK; supervision, project administration, funding acquisition, MIK.

## Funding

This research was funded by the National Institutes of Health, grant number R35 GM126944 to MIK. AES is supported by an American Cancer Society postdoctoral fellowship 721690.

## Data Availability Statement

Proteomic data sets can be found at Harvard Dataverse (https://doi.org/10.7910/DVN/IFKRBT). Genomic data sets can be found at NCBI GEO database (accession# GSE201033).

## Acknowledgments

We thank Dr. C.A. French for introducing us to NC research, and Dr. J.L. Makofske for initial work on this project. We thank Ross Tomaino (Taplin Mass Spectrometry Facility, Harvard Medical School) for expert help with analyzing the proteomics samples. We thank Drs. T. Martin and S.J. Elledge for the pAAV-tagBFP U6-gRNA expression vector. Finally, we thank H.J. Kang for helpful assistance with data submission.

## Conflicts of Interest

The authors declare no conflict of interest.

## Notes

### Competing Interest Statement

The authors have declared no competing interest.

### Summary of Updates

Text and supplemental files have been revised for clarity.

