## Supplementary material for "cMYC protein interactions and chromatin association in NUT carcinoma": Datafile S1

pAVV-MCS-NheI-5'cMyc/BamHI-ATG-loxP-Blasti-loxP-Pra-Bio-3'cMyc

gggttcctgcggccgcgctagcgccttcaggtggcgcaaaactttgtgccttggattttggcaaattgtattcctcaccgccacctcccgcggcttcttaagggcgccagggccgatttcgattcctctgccgctgcggggccgactcccgggctttgcgctccgggctcccgggggagcgggggctcggcgggcaccaagccgctggttcactaagtgcgtctccgagatagcaggggactgtccaaagggggtgaaagggtgctccctttattcccccaccaagaccacccagccgctttaggggatagctctgcaaggggagaggttcgggactgtggcgcgcactgcgcgctgcgccaggtttccgcaccaagacccctttaactcaagactgcctcccgctttgtgtgccccgctccagcagcctcccgcgacgatgataacttcgtatagcatacattatacgaagttatatgccaagcctttgtctcaagaagaatccaccctcattgaaagagcaacggctacaatcaacagcatccccatctctgaagactacagcgtcgccagcgcagctctctctagcgacggccgcatcttcactggtgtcaatgtatatcattttactgggggaccttgtgcagaactcgtggtgctgggcactgctgctgctgcggcagctggcaacctgacttgtatcgtcgcgatcggaaatgagaacaggggcatcttgagcccctgcggacggtgccgacaggtgcttctcgatctgcatcctgggatcaaagccatagtgaaggacagtgatggacagccgacggcagttgggattcgtgaattgctgccctctggttatgtgtgggagggcggatccggagccacgaacttctctctgttaaagcaagcaggagacgtggaagaaaaccccggtcccataacttcgtatagcatacattatacgaagttatatgcaggccttgcgcaacacgatgaagccgtggacaacaaattcaacaaagaacaacaaaacgcgttctatgagatcttacatttacctaacttaaacgaagaacaacgaaacgccttcatccaaagtttaaaagatgacccaagccaaagcgctaaccttttagcagaagctaaaaagctaaatgatgctcaggcgccgaaagtagacgcgaattgtgatatacctacaactgcttctctcgagagcggctcctcggctggaaaggccggtgaaggtgaaatccctgcccctcttgctggtaccgtttctaagatactggtaaaagaaggtgacactgttaaagctggtcaaacagttctggtgctggaggctatgaaaatggagacagaaattaacgctcctactgacggaaaagttgaaaaggtgttagttaaggaaagagatgctgttcaaggtggtcaaggtctaatcaagatcggcgttcgaattcggcgcgccgatggtggaggtggatctcccctcaacgttagcttcaccaacaggaactatgacctcgactacgactcggtgcagccgtatttctactgcgacgaggaggagaacttctaccagcagcagcagcagagcgagctgcagcccccggcgcccagcgaggatatctggaagaaattcgagctgctgcccaccccgcccctgtcccctagccgccgctccgggctctgctcgccctcctacgttgcggtcacacccttctcccttcggggagacaacgacggcggtggcgggagcttctccacggccgaccagctggagatggtgaccgagctgctgggaggagacatggtgaaccagagtttcatctgcgacccggacgacgagaccttcatcaaaaacatcatcatccaggactgtatgtggagcggcttctcggccgccgccaagctcgtctcagatccacgcgtgggccgcaggaacc

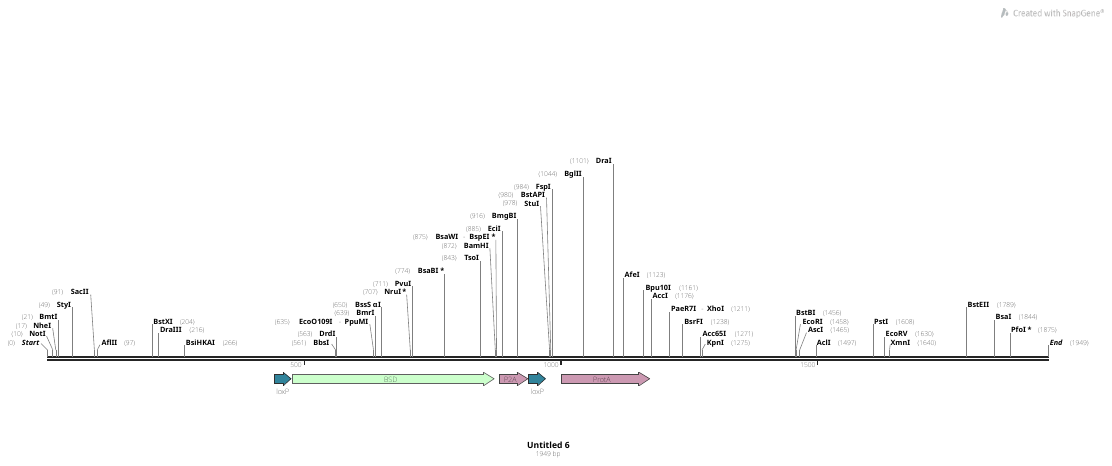

pAAV-5'-cMYC-ATG-LoxP-Blasti-loxP-N-PrA-3'-cMYC

gggttcctgcggccgcgctagcgccttcaggtggcgcaaaactttgtgccttggattttggcaaattgttttcctcaccgccacctcccgcggcttcttaagggcgccagggccgatttcgattcctctgccgctgcggggccgactcccgggctttgcgctccgggctcccgggggagcgggggctcggcgggcaccaagccgctggttcactaagtgcgtctccgagatagcaggggactgtccaaagggggtgaaagggtgctccctttattcccccaccaagaccacccagccgctttaggggatagctctgcaaggggagaggttcgggactgtggcgcgcactgcgcgctgcgccaggtttccgcaccaagacccctttaactcaagactgcctcccgctttgtgtgccccgctccagcagcctcccgcgacgatgataacttcgtatagcatacattatacgaagttatatgccaagcctttgtctcaagaagaatccaccctcattgaaagagcaacggctacaatcaacagcatccccatctctgaagactacagcgtcgccagcgcagctctctctagcgacggccgcatcttcactggtgtcaatgtatatcattttactgggggaccttgtgcagaactcgtggtgctgggcactgctgctgctgcggcagctggcaacctgacttgtatcgtcgcgatcggaaatgagaacaggggcatcttgagcccctgcggacggtgccgacaggtgcttctcgatctgcatcctgggatcaaagccatagtgaaggacagtgatggacagccgacggcagttgggattcgtgaattgctgccctctggttatgtgtgggagggcggatccggagccacgaacttctctctgttaaagcaagcaggagacgtggaagaaaaccccggtcccataacttcgtatagcatacattatacgaagttatatgcaggccttgcgcaacacgatgaagccgtggacaacaaattcaacaaagaacaacaaaacgcgttctatgagatcttacatttacctaacttaaacgaagaacaacgaaacgccttcatccaaagtttaaaagatgacccaagccaaagcgctaaccttttagcagaagctaaaaagctaaatgatgctcaggcgccgaaagtagacgcgaattgtgatatacctacaactgcttctggtggaggtggatctcccctcaacgttagcttcaccaacaggaactatgacctcgactacgactcggtgcagccgtatttctactgcgacgaggaggagaacttctaccagcagcagcagcagagcgagctgcagcccccggcgcccagcgaggatatctggaagaaattcgagctgctgcccaccccgcccctgtcccctagccgccgctccgggctctgctcgccctcctacgttgcggtcacacccttctcccttcggggagacaacgacggcggtggcgggagcttctccacggccgaccagctggagatggtgaccgagctgctgggaggagacatggtgaaccagagtttcatctgcgacccggacgacgagaccttcatcaaaaacatcatcatccaggactgtatgtggagcggcttctcggccgccgccaagctcgtctcagatccacgcgtgggccgcaggaacc

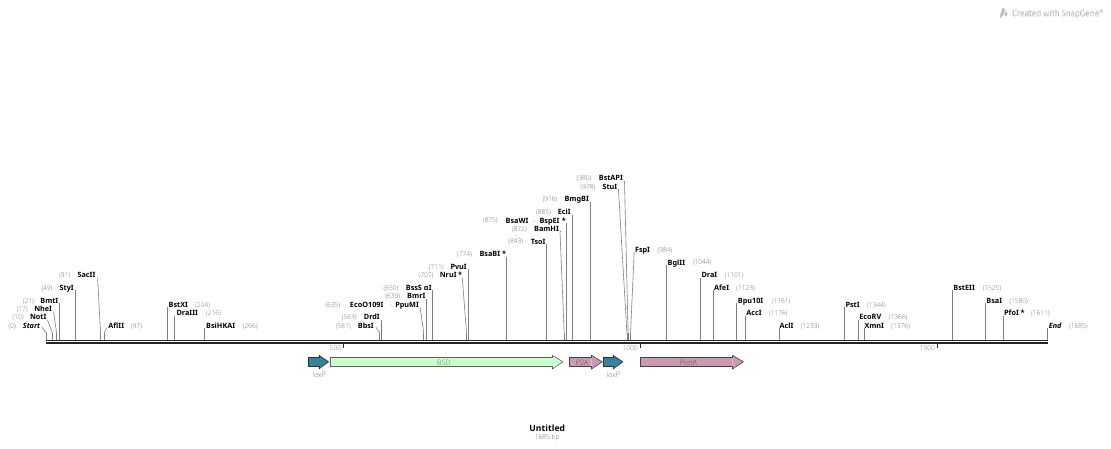
