## Supplemental Figures for "cMYC protein interactions and chromatin association in NUT carcinoma"

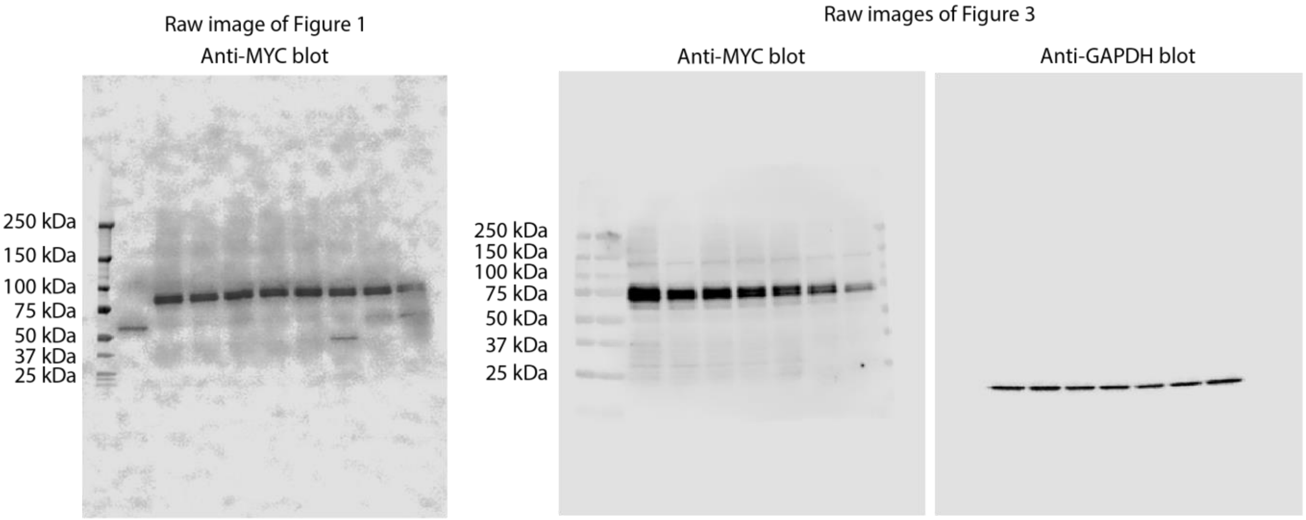


**Supplemental Figure S1.** Uncropped Western Blot images used for Figure 1 and Figure 3.

**98th percentile NC797_Myc_BioTAP1 and**
**98th percentile NC797_Myc_BioTAP2 IPs**

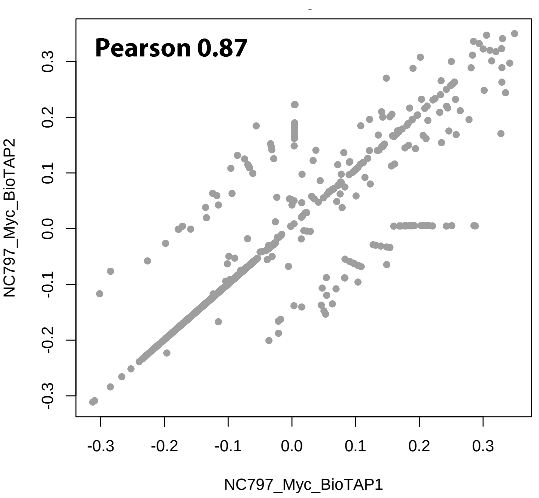


**98th percentile NC797_NPrA_Myc_noJQ1_IP_rep1 and**
**98th percentile NC797_NPrA_Myc_ noJQ1_IP_rep2 IPs**

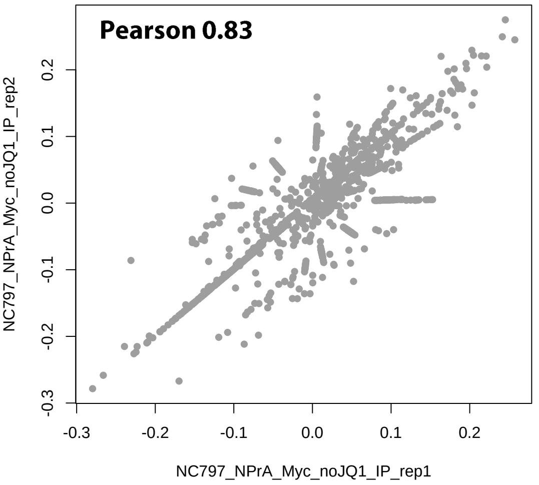


**98th percentile NC797_NPrA_Myc_12hrJQ1_IP_rep1 and**
**98th percentile NC797_NPrA_Myc_ 12hrJQ1_IP_rep2 IPs**

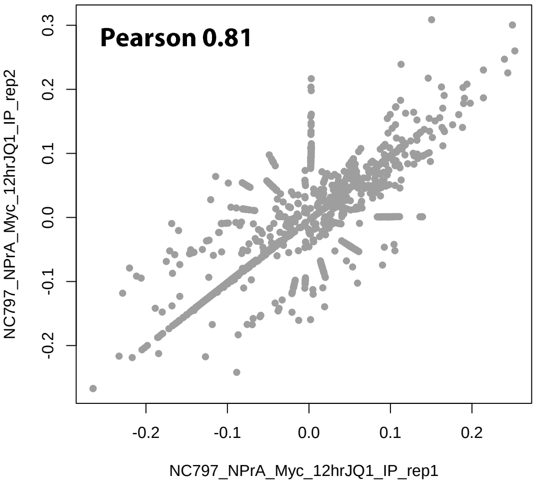


**Supplemental Figure 2.** Correlation of technical replicates of BioTAP-XL experiments, with Pearson Correlation values provided.

| | | HEK293 | | | NC797 | | |
|---|---|---|---|---|---|---|---|
| | | Input | BioTAP #1 | BioTAP #2 | Input | BioTAP #1 | BioTAP #2 |
| NuA4 | DMAP1 | 1 | 14 | 8 | 1 | 9 | 9 |
| | EP400 | 3 | 117 | 67 | 8 | 58 | 66 |
| | EPC1 | 0 | 23 | 11 | 0 | 6 | 10 |
| | EPC2 | 0 | 21 | 14 | 0 | 9 | 8 |
| | KAT5 | 0 | 20 | 10 | 1 | 10 | 8 |
| | MBTD1 | 0 | 13 | 8 | 0 | 5 | 5 |
| | TRRAP | 11 | 293 | 159 | 8 | 107 | 98 |
| | YEATS4 | 2 | 15 | 10 | 1 | 6 | 8 |
| SAGA/STAGA | ATXN7 | 0 | 10 | 5 | 0 | 0 | 1 |
| | ATXN7L2 | 0 | 7 | 2 | 0 | 0 | 0 |
| | ATXN7L3 | 0 | 4 | 3 | 0 | 0 | 0 |
| | KAT2A | 0 | 9 | 6 | 0 | 0 | 0 |
| | SUPT20H | 0 | 11 | 9 | 1 | 0 | 0 |
| | SUPT7L | 0 | 5 | 2 | 0 | 0 | 0 |
| | TADA1 | 0 | 7 | 5 | 0 | 0 | 0 |
| | TADA2B | 0 | 10 | 7 | 2 | 0 | 1 |
| | TADA3 | 0 | 10 | 5 | 1 | 0 | 0 |
| | TAF10 | 0 | 3 | 2 | 2 | 0 | 0 |
| | TAF5L | 0 | 11 | 10 | 0 | 1 | 1 |
| | TAF6L | 0 | 16 | 12 | 0 | 3 | 2 |
| | TAF9 | 2 | 13 | 11 | 1 | 1 | 4 |
| | TAF9B | 0 | 2 | 2 | 0 | 0 | 0 |

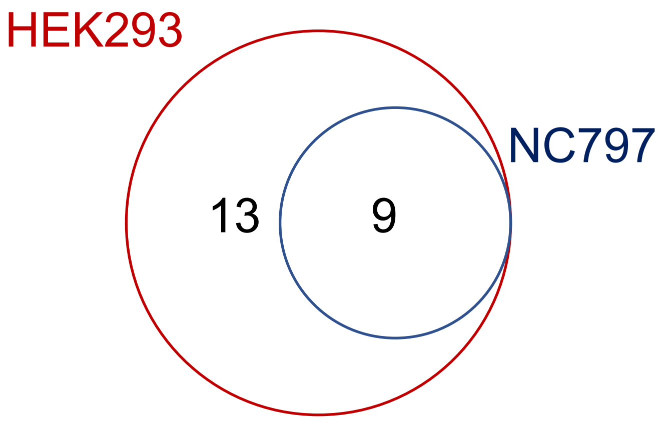


**Supplemental Figure 3.** Overlap of NuA4 and SAGA/STAGA components in MYC BioTAP experiments from HEK293 or NC797 cells.
